## Supplementary Materials for "A New Mechanism To Identify Cost Savings in NHS Prescribing: Minimising “Price-Per-Unit”"

### Appendix B – BNF codes excluded from price-per-unit analysis

0302000C0____BE
0302000C0____BF
0302000C0____BH
0302000C0____BG
0904010H0%
0904010H0%
1311070S0____AA
1311020L0____BS
0301020S0____AA
190700000BBCJA0
0604011L0BGAAAH
1502010J0____BY
0107010S0AAAGAG
060106000BBAAA0
190201000AABJBJ
190201000AABKBK
190201000AABLBL
190201000AABMBM
190201000AABNBN
190202000AAADAD
